# Metadata-Guided Visual Representation Learning for Biomedical Images

**DOI:** 10.1101/725754

**Authors:** Stephan Spiegel, Imtiaz Hossain, Christopher Ball, Xian Zhang

## Abstract

**Motivation:** The clustering of biomedical images according to their phenotype is an important step in early drug discovery. Modern high-content-screening devices easily produce thousands of cell images, but the resulting data is usually unlabelled and it requires extra effort to construct a visual representation that supports the grouping according to the presented morphological characteristics.

**Results:** We introduce a novel approach to visual representation learning that is guided by metadata. In high-context-screening, meta-data can typically be derived from the experimental layout, which links each cell image of a particular assay to the tested chemical compound and corresponding compound concentration. In general, there exists a one-to-many relationship between phenotype and compound, since various molecules and different dosage can lead to one and the same alterations in biological cells.

Our empirical results show that metadata-guided visual representation learning is an effective approach for clustering biomedical images. We have evaluated our proposed approach on both benchmark and real-world biological data. Furthermore, we have juxtaposed implicit and explicit learning techniques, where both loss function and batch construction differ. Our experiments demonstrate that metadata-guided visual representation learning is able to identify commonalities and distinguish differences in visual appearance that lead to meaningful clusters, even without image-level annotations.

**Note:** Please refer to the supplementary material for implementation details on metadata-guided visual representation learning strategies.

## 1 Introduction and Background

High-content screening aims at automating discovery in cell biology and drug development using large amounts of microscopy images [20]. Most imaging assays are designed to study the effect of environmental perturbations on cell lines, but the resulting cellular phenotypes are typically unknown and too subtle to identify manually, leading to unlabelled datasets [11]. In order to infer the relationship between environmental perturbations and phenotypic characteristics, which is of high importance for drug target validation as well as the identification of new lead compounds [26], the unlabelled images have to be clustered according their visual appearance.

There exists a plethora of clustering algorithms for (biomedical) images, which employ unsupervised learning techniques to derive insights from the data itself and to make data-driven decisions without external bias [25]. More recently, unsupervised representation learning has been employed to identify novel cellular phenotypes [6], detect abnormal cell morphologies [23] and analyse biomedical images in an exploratory manner [14]. However, unsupervised machine learning generally faces the challenge that it is rather difficult to evaluate whether the algorithm has learned anything meaningful about the internal data structure, since there is no objective performance measure to guide the learning [25].

Hence, we propose metadata-guided visual representation learning, which exploits contextual information to direct unsupervised learning. In case of high-content screening assays, our approach utilizes information about the tested compounds and their applied concentrations in order to facilitate learning of the intrinsic image structure. In general, our approach is able to employ any kind of metadata that allows to determine pseudo-classes, insofar as every pseudo-classes only belongs to one particular super-class of our final clustering, refer to Figure 1 and Section 1.1.

**Figure 1:**
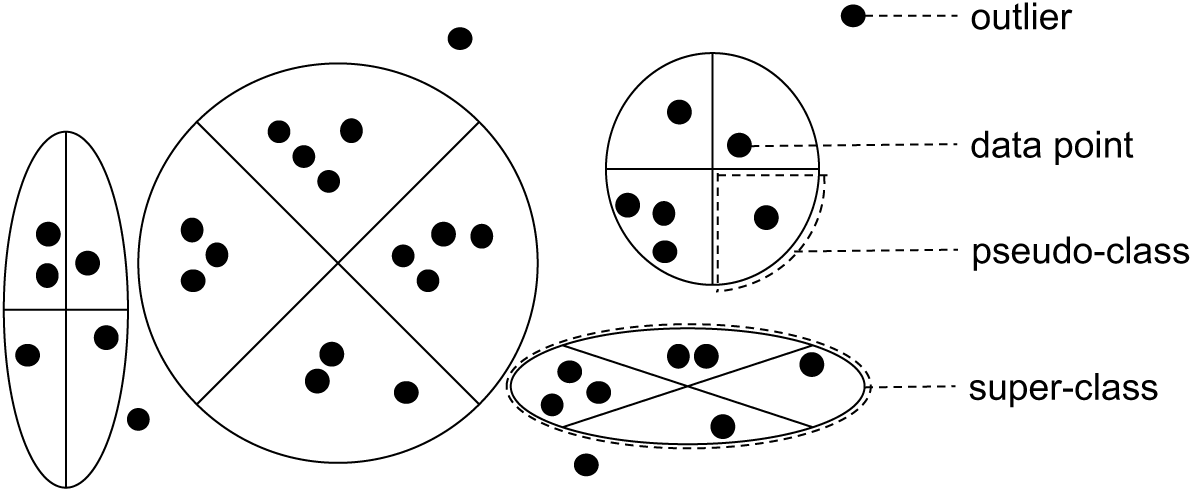
Embedded feature space, illustrating an exemplary clustering of data points into super-classes, which have been derived from pseudo-classes by means of our proposed metadata-guided visual representation learning. Please note that each ‘pseudo’-class only belongs to one particular super-class, although a super-class typically consists of multiple ‘pseudo’-classes.

We evaluate our metadata-guided visual representation learning using two opposing strategies, namely implicit learning and explicit learning. More precisely, implicit learning typically solves a pseudo task, such as training a convolutional neural network for image classification using categorical cross-entropy as loss function, to actually learn a visual representation, which is ultimately given by an intermediate layer of the respective deep learning architecture [10]. In contrast, explicit learning trains the visual representation by directly imposing constraints on the embedded feature space employing a ‘geometric’ loss function, which defines the spacial relationship between tuplets [5, 7], triplets [9, 19, 27] or, as in our study, ‘N’-lets [22].

We assess the performance of our metadata-guided visual representation learning approach on two different datasets. First, we test on a MNIST [12], a benchmark dataset that is well-known inside of the computer vision and deep learning community. Second, we evaluate on BBBC021 [3, 13], a real-world biomedical image dataset that is widely-used in the field of computational biology. Since both of the selected datasets are relatively small in comparision to the number of trainable parameters found in modern deep learning architectures, we decided to employ transfer learning [2, 17], where only the last couple of neural network layers are readjusted to fit the new domain. For both implicit and explicit learning strategy, we decided to employ the intensively-studied VGG16 model [21] that was pretrained on the ImageNet dataset [18].

### 1.1 Formal Problem Definition

In supervised learning we usually have a set of *m* training examples of the form *x* = {(*x*^(1)^, *y*^(1)^), …, (*x*^(*m*)^, *y*^(*m*)^)}, with *x* ∈ ℝ^#*features*^ and *y* ∈ ℝ^#*classes*^, such that *x*^(*i*)^ is the feature vector of the *i*-th example and *y*^(*i*)^ is the given class label. A machine learning algorithm seeks a function *g*:*X*→*Ŷ*, where X is the input space and *Ŷ* is the output space. Typically we want the probability for a predicted class label to be close to the actual class label, i.e. *ŷ*^(*i*)^ ≈ *y*^(*i*)^. The cost *𝒥* of a parametrized model, with weight matrix *w* and bias term *b*, is defined as 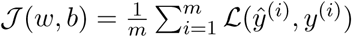, where the loss function ℒ penalizes deviations in predicted and actual class membership.

In case of metadata-guided visual representation learning, we assume a set of *m* training images of the form *x* = *{*(*x*^(1)^, *p*^(1)^), …, (*x*^(*m*)^, *p*^(*m*)^)*}*, with *x* ∈ℝ^#*pixel*^ and *p* ∈ ℝ^#*pseudo-classes*^, where *p*^(*i*)^ is a ‘pseudo’-label that we consider as a proxy to infer *ŷ*^(*i*)^, which in turn allows us group the images according to the estimated class membership *Ŷ* without ever knowing the actual super-classes *Y*, a.k.a. clustering or unsupervised learning. Typically the metadata subdivides our training set into many more ‘pseudo’-classes than actual ‘super’-classes, i.e. #*pseudo-classes ≫* #(*super-*)*classes*. Our approach aims at utilizing image- as well as meta-data in order to learn a function that is able to predict the (super-)class membership: *f* :*X*×*P* →*Ŷ*.

## 2 Model Architecture and Learning Strategy

Figure 2 illustrates the deep learning architecture that we have employed for visual representation learning and subsequent image clustering. Our architecture builds on top off the well-known VGG16 model [21], which has been pre-trained on the ImageNet dataset [18], consisting of 1M pictures from 1K categories, like cats and dogs. Although we aim at investigating images from a rather different domain, the first couple of VGG16’s building blocks (layer 0-14) have been pre-trained to extract low-level image features, such as edges and textures, which represent a quintessential prerequisite for solving almost any image analysis task. Since we are not interested in classifying cats and dogs, we remove VGG16’s top layers and add another building block, which is used to learn visual representations or embeddings that potentially help us to group biomedical images.

**Figure 2:**
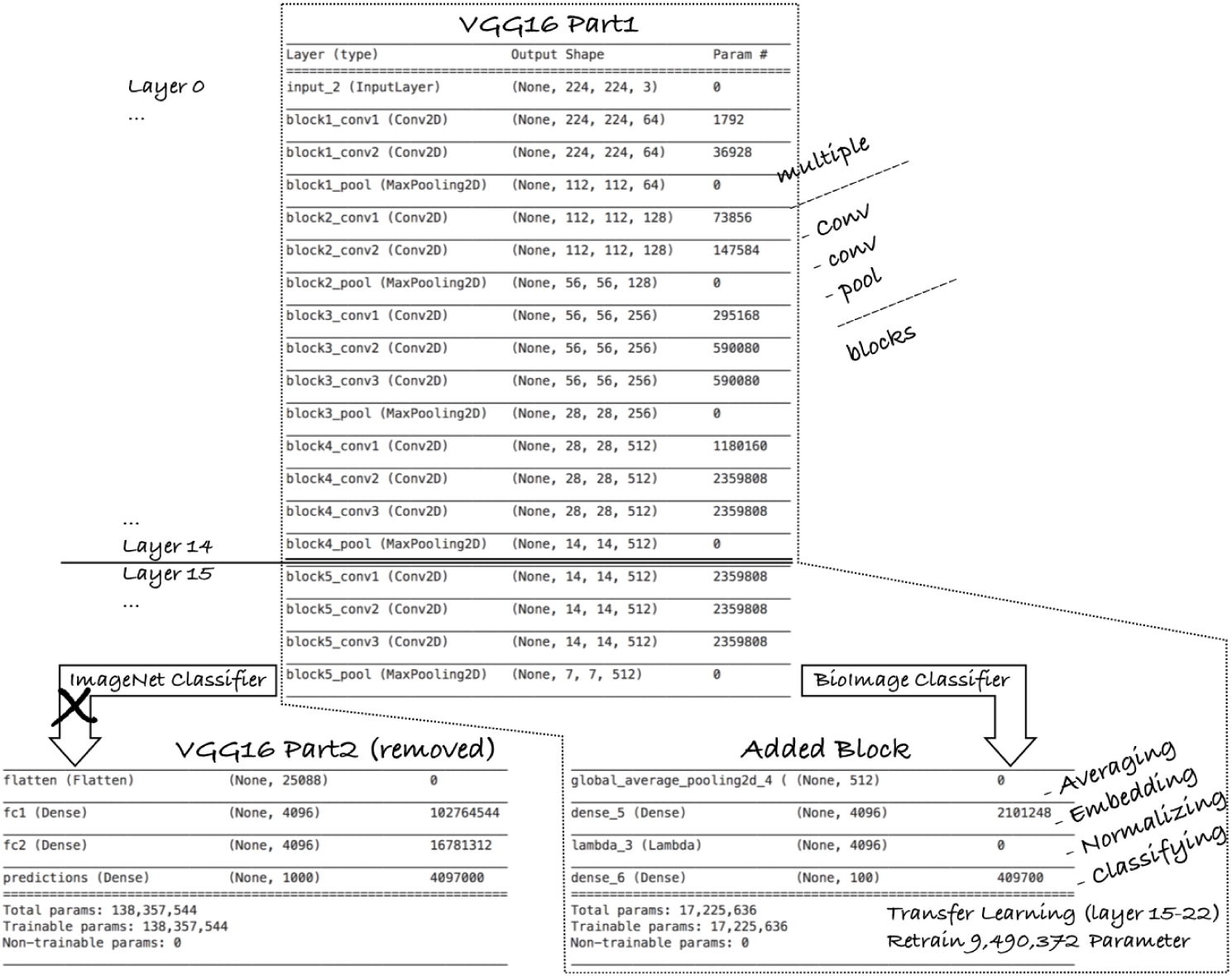
Our proposed visual representation learning architecture, which reuses multiple pre-trained VGG16 blocks and appends additional layers for averaging, embedding, normalization and classification. We merely re-train layer 15-22 (about 9M parameters) in the transfer learning step.

In the so-called transfer learning step, we adjust our assembled model to the biomedical domain, re-training the parameters of VGG16’s last building block and our newly added building block, e.g. on cell images captured by a high-content-screening device. By only re-training the last part of our deep learning model (layer 15-22), we decrease the number of parameters (from roughly 138M to about 9M), leading to reduced computational demand and less training time. In other words, we get some powerful low-level feature extractors for free and just need to take care about the high-level visual representations or embeddings.

The building block that we have added to the VGG16 model, as shown in Figure 2, consists of multiple layers that: (i) <u>aggregate</u> the features extracted by the previous building block, (ii) <u>embed</u> the aggregated features in a lower-dimensional space, (iii) <u>normalize</u> the embedding to facilitate distance measures, and ultimately (iv) <u>classify</u> the input images. In our above illustrated example, the output layer aims at classifying the input images into 100 categories, however the number of classes depend on the problem/dataset at hand.

Given the described architecture, we have investigated two different ways of visual representation learning, namely *implicit* and *explicit*. The following subsections explain the main differences.

### 2.1 Implicit Representation Learning

We can learn a visual representation *implicitly* by actually training a deep neural network to classify images, as performed by our architecture shown in Figure 2. Although we mainly optimize the classification accuracy, we *implicitly* adjust the parameters of the intermediate embedding layer, which gives us a low-dimensional feature representation of our input images.

Assuming a dataset with *n* classes, our output layer consists of *n* softmax units, one for each category. Given *m* samples, the model parameters (weight matrix *w* and bias *b*) are optimized according to the summed loss of the categorical cross-entropy:

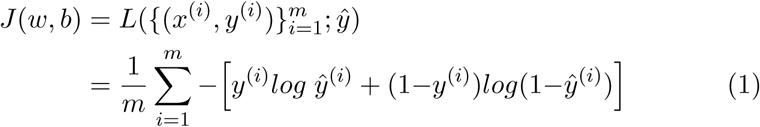

where (*x*^(*i*)^, *y*^(*i*)^) is the *i*-th tuple of (input image, actual class label) and *ŷ*^(*i*)^ is the predicted class label that our model has inferred from the processed input image *x*^(*i*)^. The cost function *𝒥* accounts for the loss of all *m* samples, whereas the loss *L* for each individual sample is computed by comparing the actual class label *y*^(*i*)^ with the predicted class label *ŷ*^(*i*)^.

Typically we train a deep neural network iteratively by computing the cost of a mini-batch and, subsequently, updating the model parameters, e.g., by means of stochastic gradient descent. The back-propagation algorithm, as the name suggests, is used to propagate the error back through all layers, including our embedding layer. We hypothesize that the *implicitly* trained embedding layer provides us with meaningful visual representations as the classification error is minimized. Evidence is presented in Section 4.

### 2.2 Explicit Representation Learning

We can learn a visual representation *explicitly* by removing the classification layer and using the normalized embedding layer as model output, please refer to Figure 2. Since the normalized embedding layer does not allow us to predict class label, we need to use an alternative cost function that guides parameter learning in the right direction.

In this study we investigate the mutli-class n-pair loss function, for short ‘n-let’ loss, that was originally proposed by *K. Sohn* [22]. Same as with contrastive loss [5, 7] and triplet loss [9, 19, 27], the main idea is that we project our input images into a lower-dimensional space that clusters same-class images and separates different-class images.

Given a mini-batch of *m* pairs, the ‘n-let’ loss aims at pulling together same-class instances from one pair, while simultaneously pushing away all different-class instances from other pairs:

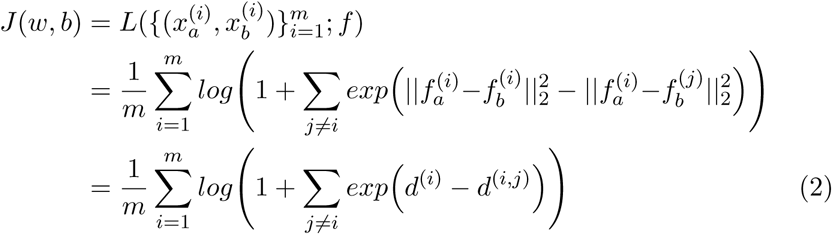

where 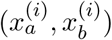 is a tuple of same-class instances and 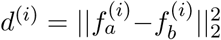 is the distance between their normalized embeddings that our models has computed by processing the respective input images. Conversely, the dissimilarity between different-class images is expressed as 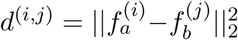.

In general, we want the similarity between same-class images to be high and the distance between different-class images to be large. To paraphrase, we aim to minimize intra-class similarity and maximize inter-class distance simultaneously. This is reflected in the above cost function (2), since the loss converges to zero as the logarithmic function comes closer to one, which is the case if the exponential function moves towards the negative numbers, meaning that the distance between same-class images is significantly smaller than the distance between different-class images, i.e. *d*^(*i*)^ ≪ *d*^(*i,j*)^.

A thorough experimental comparison of *implicit* and *explicit* representation learning is presented in Section 4. Prior to this, we are going to explain the use of metadata in the following Section 3

## 3 Image Data and Metadata

In visual representation learning, we aim at building robust machine learning models that understand and distinguish high-level visual concepts. Even though images live in high-dimensional spaces and, hence, are intrinsicly very rich in information, many image datasets come <u>without</u> concept label, because annotation is often time/cost-intensive [8, 24].

Missing label prevent us from applying supervised learning, which maps images to known visual concepts, and confine ourselves to <u>un</u>supervised learning, which groups the <u>un</u>labeled images on the basis of underlying structural features, representing an inherently difficult and, furthermore, mathematically ill-defined task [25]. To this end, we propose to employ metadata, whenever available, to guide visual representation learning in a ‘pseudo’-supervised learning approach, where ‘pseudo’-label are derived from the metadata itself.

In general, metadata is defined as any kind of additional information that includes relevant details about the image itself as well as its production. For instance in high-content-screening, biomedical images are often tagged with their plate location (row, column, field) and, furthermore, carry information about the compound-concentration that was applied to the biological cells in the corresponding well [3, 13].

Although metadata does <u>not</u> correspond to the actual class label of an image, it can be considered as a ‘pseudo’-label that describes a <u>sub</u>group. For example, multiple compounds in different concentrations can lead to one and the same morphological characteristics, where each compound-concentration just describes a <u>sub</u>-population of the actual phenotype. Nonetheless, this contextual information can be extremely valuable, as we will demonstrated by our empirical results in Section 4.

In our experiments we consider two image datasets, one for the purpose of benchmarking and the other for providing real-world evidence.

### 3.1 Benchmark Data

MNIST [12] is one of the most popular and widely used datasets for bench-marking image analysis and processing tools. It consists of 60K plus 10K handwritten digits, which are used for training and validation respectively. The digits range from zero to nine and are grouped into ten categories. All images are gray-scale, centered and scaled to 28×28 pixel. Some random samples of the original MNIST dataset are shown in Figure 3.

**Figure 3:**
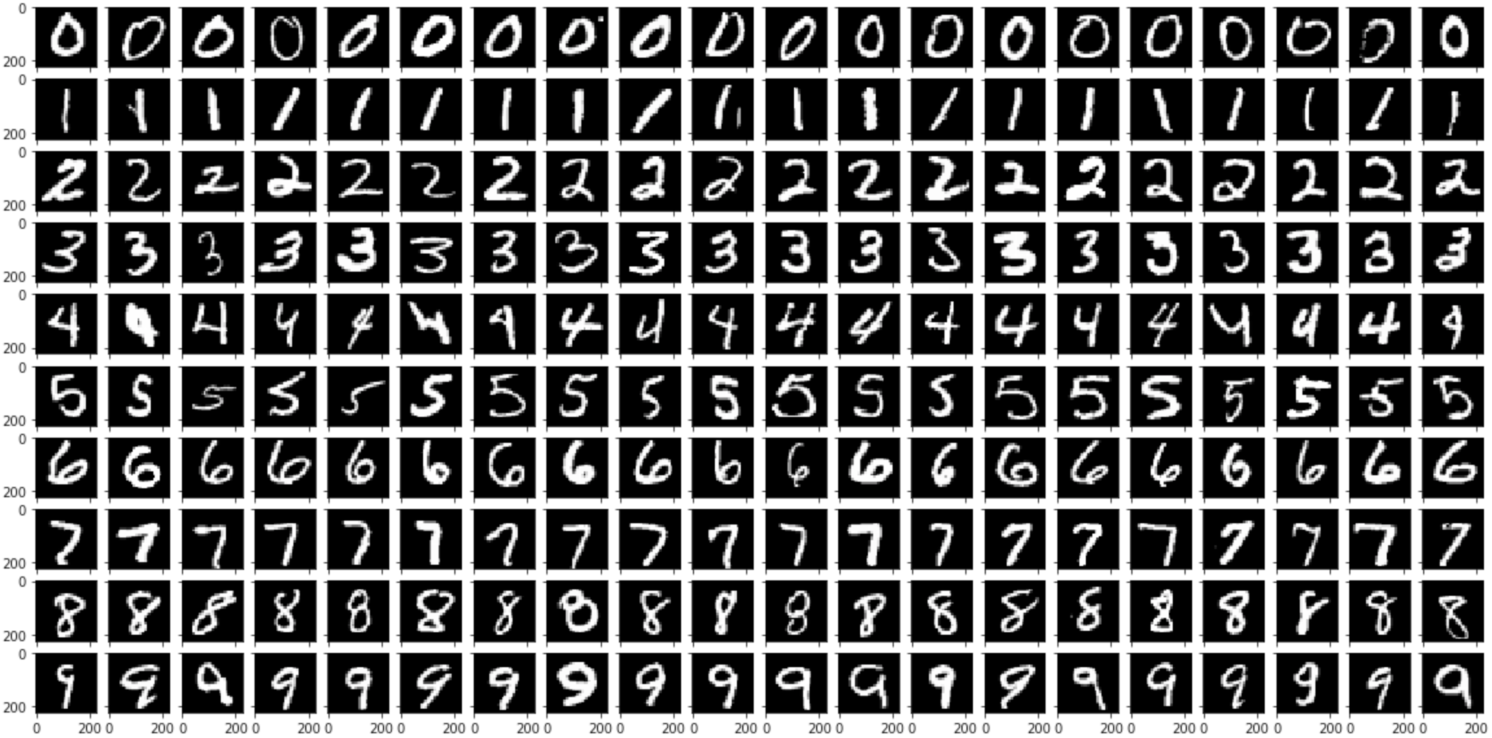
MNIST digits, each row shows one of 10 different classes and each column presents a random sample, illustrating the variance within each class.

For our experiments we need to prepare/modify the MNIST dataset in a certain way. As shown in Figure 2, our model builds on top of the VGG16 architecture[21], which only accepts input images with a certain dimension. Hence, we rescale and triplicate the gray-scale channel of all MNIST digits, converting 28×28×1 into 224×224×3 dimensional images.

Furthermore, we divide the original categories into multiple subclasses, which can be considered as metadata or pseudo-label that we subsequently exploit for representation learning. For example, we can generate 100 pseudo-label by splitting each of the 10 original categories into 10 random sub-populations. In general, the following assumptions should hold true:

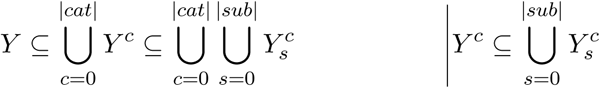

where *Y* is the set of all labelled instances, *Y*^*c*^ is the set of all instances labelled as category *c*, and 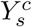 is the set of all category *c* instances that have been randomly assigned to subclass *s*.

To draw an analogy to our previous biomedical imaging example, we can consider individual subclasses as different compounds and categories as cell phenotypes. Typically there exists a many-to-one relationship between compound and phenotype. In our setting, we are trying to group images with same phenotype by utilizing compound information, which is in fact just a pseudo-label or rather metadata.

By dividing the MNIST categories into an arbitrary number of sub-classes, we can measure the performance of our previously proposed visual representation learning techniques. Usually the number of subclasses should be bigger than one and smaller than the number of samples in a category, since one subclass per sample would set us back to unsupervised learning and one subclass per category basically equates to supervised learning, which assumes labelled data that we are missing. The critical role of the subclass size is emphasized in our empirical study in Section 4

### 3.2 Real-World Data

In order to demonstrate that our proposed visual representation learning techniques are also applicable to real-world datasets, we furthermore provide experimental results on the image set BBBC021 [3], available from the Broad Bioimage Benchmark Collection [13].

BBBC021 images are of MCF-7 breast cancer cells treated for 24 h with a collection of 103 compound-concentrations (38 compounds at 1-7 concentrations each). The tested compound-concentrations have been identified to cause one of 12 primary mechanisms of action, where only 6 mechanisms have been confirmed visually (due to subtle phenotypical differences) and the remaining 6 mechanisms have been defined based on the literature. The cells were fixed, labeled for DNA, Actin, and Tubulin, and imaged by fluorescent microscopy. There are 39,600 image files (13,200 fields of view imaged in three channels – one for DNA, F-Actin and B-Tubulin) in TIFF format. This image set provides a basis for testing image-based profiling methods wrt. to their ability to predict the mechanisms of action of a compendium of drugs [3, 13].

For our study we have prepared the BBBC021 dataset in the following way. First of all we have used CellProfiler [4] to identify the centroid of each individual cell in all of the 13,200 images. Subsequently we have filtered all images that show less than 20 cells, where cells that extend beyond the border of an image are excluded from counting and further analysis. From the remaining ∼12, 500 images we select only those that have been identified to show one of 12 phenotypes or rather mechanisms of action, leaving us with exactly 3,011 images. For each of these images we have randomly choosen 22 cells and cropped a 224×224 region around the centroids identified previously. Since we have considered all of the three channels, we obtain 224×224×3 dimensional tensors that fits the input size of our deep neural network, shown in Figure 2. Consequently, the final dataset that we have distilled has shape (3011, 22, 224, 224, 3) and is available on request.

The selected 3,011 images illustrate 11 phenotypes that were caused by 57 compound-concentrations. We have performed a 90%/10% split for training/validation, such that both sets contains examples of all 57 tested compound-concentrations, producing tensor shapes (2719, 22, 224, 224, 3) and (292, 22, 224, 224, 3). For model training we have employed all 57 tested compound-concentrations as label, whereas for validation we have just considering the 11 phenotypes, resulting in tensor shapes (2719, 57) and (292, 11) respectively.

This setup allows us to evaluate if our model has been able to recover some of the morphological characteristics by merely using information about the compound-concentrations. In that sense, compound-concentrations can be considered as pseudo-label or metadata, which are used to guide visual representation learning, where the embedding is trained in a way that it allows to group/cluster the individual phenotypes or mechanisms of actions.

We present empirical results for both implicit and explicit representation learning on our prepared MNIST benchmark dataset as well as on the real-world BBBC021 dataset in the following Section 4.

## 4 Empirical Results

In our empirical study we have evaluated implicit and explicit representation learning, as discussed in Section 2, on both benchmark and real-world data, as described in Section 3. The validation of the learned representations is presented in Subsection 4.1. In addition, Subsection 4.2 includes a close inspection of the established clustering. However, the discussion of all results is kept separately (on intention) and placed in Section 5.

There exist multiple model parameters that potentially influence learning and, therefore, the resulting visual representations. In our study, we merely vary the number of ‘pseudo’-classes that guide our learning algorithm and fix all other parameters, since the explorable space is quite large. More precisely, we employ stochastic gradient descent with *learning rate* = 0.005 as well as *momentum* = 0.9, and set *embedding dimension* = 4096. Results are presented for training over 1 to 5 epochs.

Note that the size of the output layer for the implicit model depends on the number of ‘pseudo’-classes provided by the metadata, see Figure 2. Moreover, we only re-train weights for layer 15 and deeper, using categorical cross-entropy for implicit learning and ‘n-let’ loss for explicit learning, refer to Section 2.1 and 2.2 respectively.

Before we dive into the empirical results, we want to explain how to even assess performance for an established clustering. Since we know the actual class label for both training and validation set, we can use those to define an external evaluation metric. More precisely, after our model has learned an embedding from the training set (in an unsupervised fashion), we can measure the distance between a pair of embedded validation images in order to decide whether the two data points belong to the same class or not, and ultimately compare this decision with the ground truth. In our study, we report the average distance for same-class, different-class, and any-class pairs. Based on the pair-wise distances and a predefined threshold, e.g. cutoff at 1.0 distance, we are able to make same-class/different-class decisions and, given the ground truth, estimate the accuracy of our decision making. For our experiments, we report same-class, different-class, and any-class/overall accuracy.

### 4.1 Cluster Validation

#### 4.1.1 Benchmark Validation

In order to make a fair comparison between implicit and explicit visual representation learning, we need to train both models on the same amount of images. For implicit learning we usually specify the number of epochs, which reflect the number of times the model is seeing all training instances. In contrast, for explicit learning we just define the batch size, which is selected based on the number of ‘pseudo’-classes provided by the available metadata. Given *n*^2^ ‘pseudo’-classes, we typically construct batches of size 2 × *n*, containing *n* same-class image pairs utilized by the ‘n-let’ loss. Note that *n* is typically an estimate of the number of super-classes (e.g. number of MNIST digits or BBBC021 phenotypes) that we expect/aim to recover from the unlabelled dataset.

For example, in case of our MNIST benchmark dataset, the implicit learning model has been trained over 5 epochs on 90% of the 60K training instances, adding up to 5 × 0.9 × 60, 000 = 270, 000 images that were utilized to tune the parameters of our deep neural network. Assuming 100 ‘pseudo’-classes and 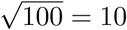 super-classes (or potential digits), we can make a fair performance comparison with our explicit learning model by constructing *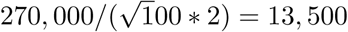*, 500 batches with 10 image pairs (*2), please refer to ‘n-let’ batch construction for explicit learning in Section 2.2.

In Table 1 we compare the performance difference between implicit and explicit representation learning for a varying number of ‘pseudo’-classes. Note that with a growing number of ‘pseudo’-classes the number of samples per ‘pseudo’-class is decreasing. In our evaluation we measure the average accuracy of predicting correctly that two test images are of same or different class, where the overall classification performance is given by their mean with respect to the number of same and different pairs in the validation set.

**Table 1:**
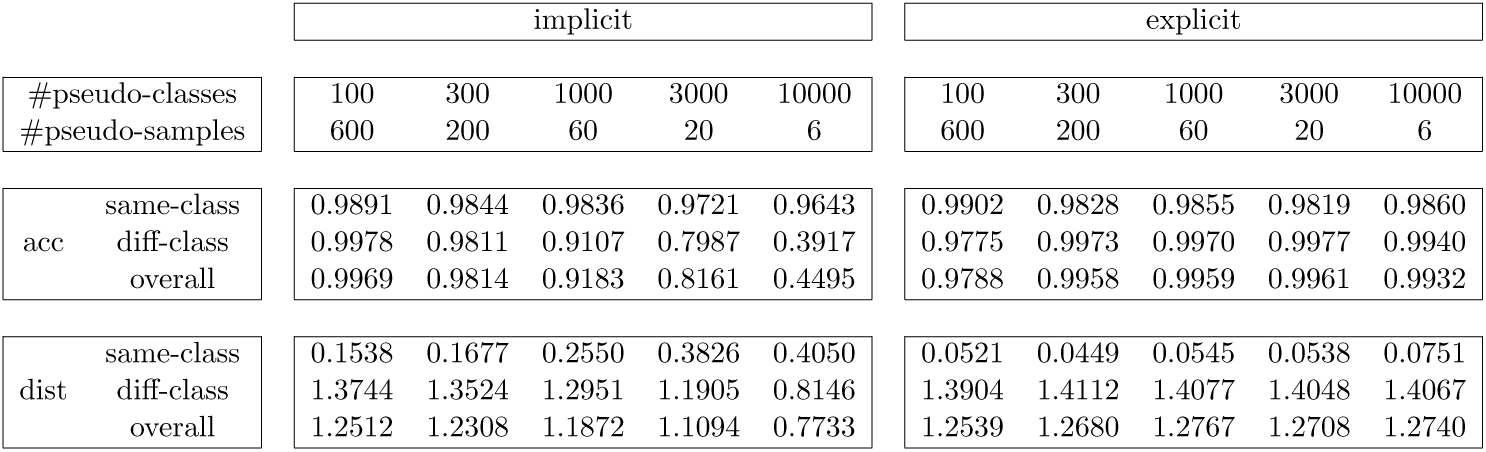
Average accuracy (acc) and average distance (dist) for implicit and explicit learning on prepared/subdivided MNIST dataset (see Section 3.1), with varying number of pseudo-classes or rather pseudo-samples per class. The numbers correspond to model performance after 5 epochs of training.

In Table 1 we furthermore juxtapose the average intra-class, inter-class and overall distance between test images in the embedded space that has been learned either implicitly or explicitly. In general, we aim to learn an embedding that produces small intra-class and high inter-class distances, such that individual super-classes (MNIST digits or BBBC021 phenotypes) are separated according their visual characteristics.

Figure 5 provides a visual comparison of implicit and explicit learning, which clearly shows the performance discrepancy in our MNIST benchmark. We plot overall accuracy and overall distance results. Please note that we have added an additional data point for 30 pseudo-classes only.

**Figure 4:**
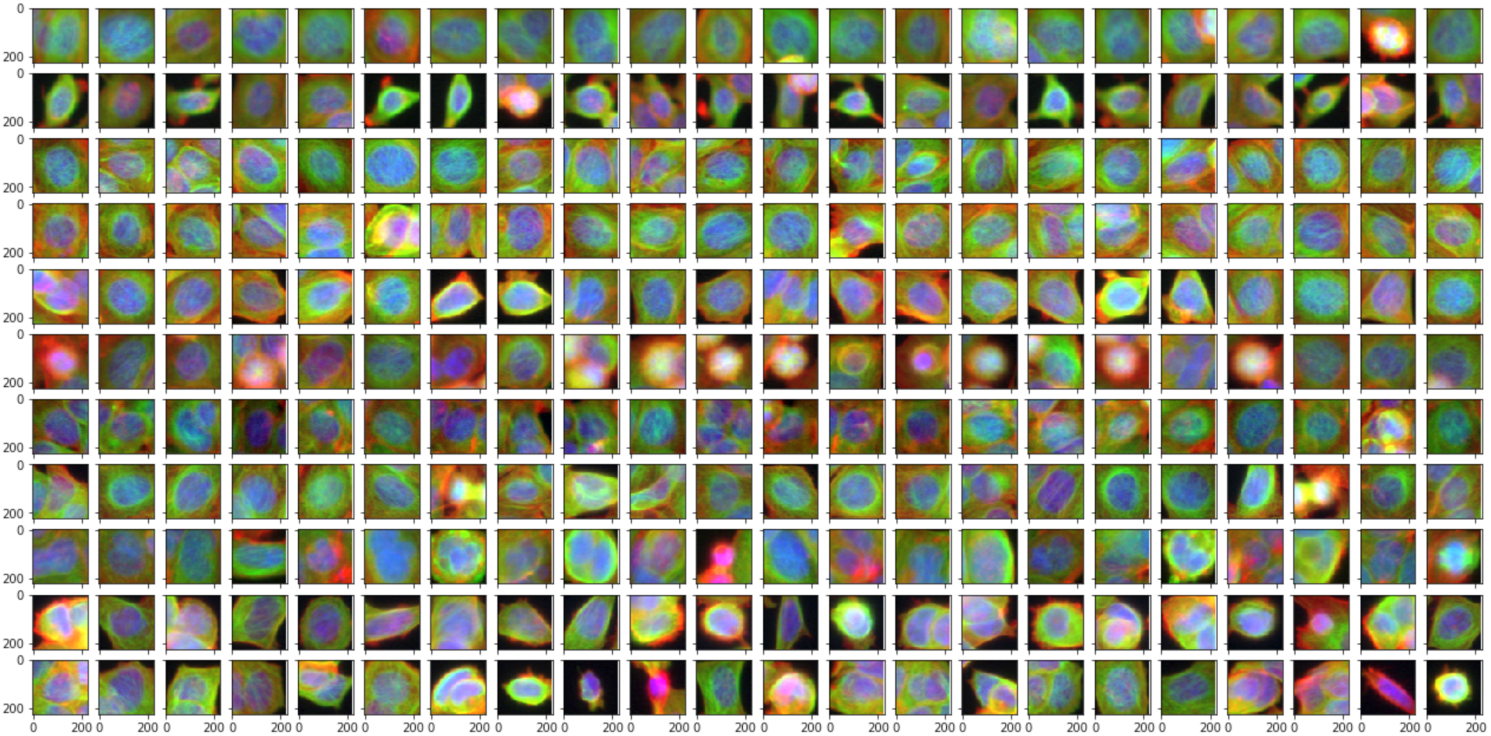
MCF-7 breast cancer cells cropped from the original BBBC021 images. Each row shows one of 11 different phenotypes, where 22 cells were randomly sampled from one and the same image. This illustrates the variance of morphological characteristics in cellular images. Furthermore, the plot demonstrates that cells with similar visual appearance are found within different phenotypes (or rather rows). High intra-class variance and high inter-class similarity present a challenge for our representation learning.

**Figure 5:**
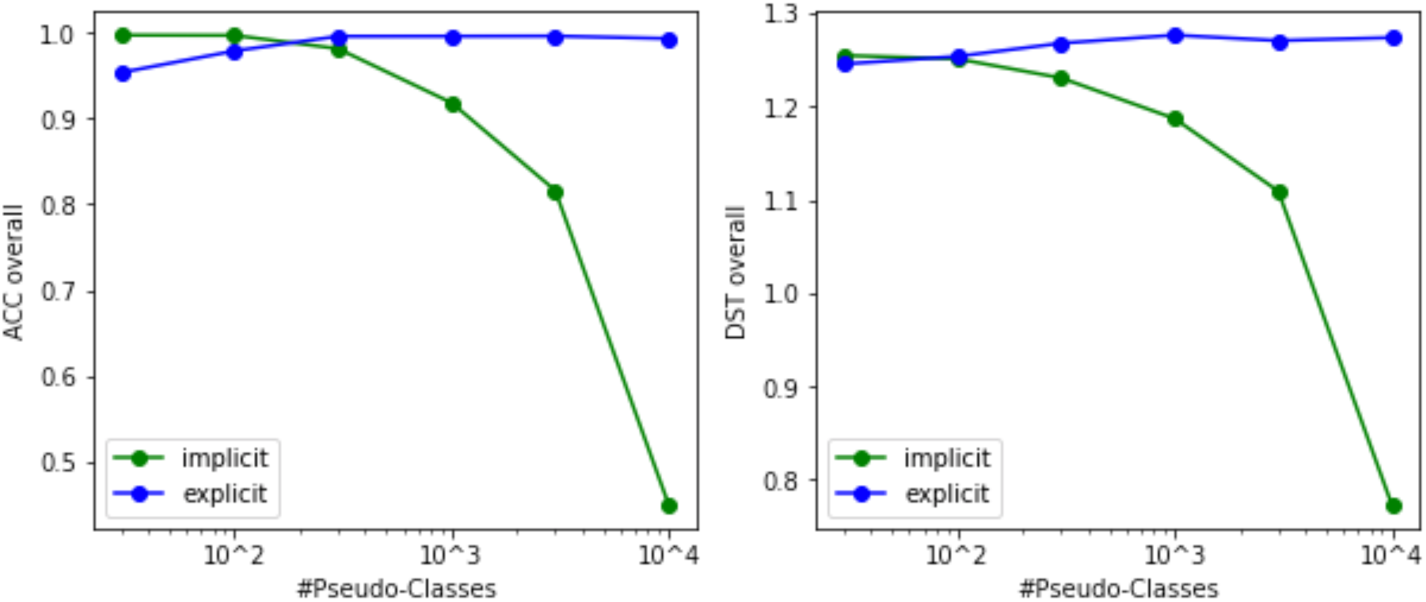
Visual comparison of implicit (green) and explicit (blue) learning approach, showing overall accuracy (left) and overall distance (right) on MNIST benchmark for varying number of ‘pseudo’-classes, refer to Table 1.

In addition, we have performed another series of benchmark experiments, where we measure model performance after a varying number of training epochs. These experiments are insightful, since the results show how the intra-class and inter-class distances evolve with more batches seen/processed by the deep neural network. We compare model training of implicit and explicit learning for 10^2^ and 10^4^ ‘pseudo’-classes in Table 2.

**Table 2:** Average accuracy (acc) and average distance (dist) for implicit and explicit learning on prepared/subdivided MNIST dataset (see Section 3.1), for varying number of epochs/batches. The training loss shows how fast our model learns from the processed data and, furthermore, indicates how well the learned embedding is going to group same-class images and separate different-class images for our validation set, as presented in the rows below.

| 10<sup>2</sup> pseudo-classes |  | implicit |  |  |  |  | explicit |  |  |  |  |
| --- | --- | --- | --- | --- | --- | --- | --- | --- | --- | --- | --- |
| #epoch<br>#batch |  | 1 | 2 | 3 | 4 | 5 | 1 | 2 | 3 | 4 | 5 |
|  |  | 2700 | 5400 | 8100 | 10800 | 13500 | 2700 | 5400 | 8100 | 10800 | 13500 |
| loss | training | 4.3547 | 3.6515 | 3.1376 | 2.8424 | 2.6747 | 1.6118 | 1.52347 | 1.48259 | 1.45858 | 1.4440 |
| acc | same-class | 0.9648 | 0.9759 | 0.9893 | 0.9910 | 0.9921 | 0.9845 | 0.9742 | 0.9728 | 0.9839 | 0.9794 |
|  | diff-class | 0.9297 | 0.9878 | 0.9938 | 0.9958 | 0.9966 | 0.9713 | 0.9959 | 0.9879 | 0.9959 | 0.9970 |
|  | overall | 0.9333 | 0.9866 | 0.9933 | 0.9953 | 0.9961 | 0.9727 | 0.9937 | 0.9865 | 0.9947 | 0.9952 |
| dist | same-class | 0.3451 | 0.2194 | 0.1802 | 0.1564 | 0.1406 | 0.1276 | 0.1007 | 0.1188 | 0.0551 | 0.0500 |
|  | diff-class | 1.2848 | 1.3432 | 1.3577 | 1.3660 | 1.3705 | 1.3802 | 1.4082 | 1.3974 | 1.4097 | 1.4111 |
|  | overall | 1.1887 | 1.2312 | 1.2369 | 1.2447 | 1.2458 | 1.2508 | 1.2763 | 1.2752 | 1.2751 | 1.2724 |

Table 2: Average accuracy (acc) and average distance (dist) for implicit and explicit learning on prepared/subdivided MNIST dataset (see Section 3.1), for varying number of epochs/batches.
| 10<sup>4</sup> pseudo-classes |  | implicit |  |  |  |  | explicit |  |  |  |  |
| --- | --- | --- | --- | --- | --- | --- | --- | --- | --- | --- | --- |
| #epoch<br>#batch |  | 1 | 2 | 3 | 4 | 5 | 1 | 2 | 3 | 4 | 5 |
|  |  | 2700 | 5400 | 8100 | 10800 | 13500 | 2700 | 5400 | 8100 | 10800 | 13500 |
| loss | training | 9.2129 | 9.2120 | 9.2111 | 9.2096 | 9.2063 | 1.6167 | 1.5272 | 1.4878 | 1.4619 | 1.4447 |
| acc | same-class | 1.0000 | 1.0000 | 1.0000 | 0.9991 | 0.9724 | 0.9511 | 0.9728 | 0.9852 | 0.9884 | 0.9800 |
|  | diff-class | 0.0000 | 0.0000 | 0.0000 | 0.0973 | 0.3464 | 0.9532 | 0.9720 | 0.9948 | 0.9979 | 0.9960 |
|  | overall | 0.0959 | 0.1027 | 0.0999 | 0.1925 | 0.4121 | 0.9530 | 0.9721 | 0.9938 | 0.9969 | 0.9944 |
| dist | same-class | 0.0808 | 0.0990 | 0.1671 | 0.2984 | 0.4259 | 0.1530 | 0.1036 | 0.0764 | 0.0481 | 0.0596 |
|  | diff-class | 0.1036 | 0.1376 | 0.2645 | 0.5284 | 0.7971 | 1.3674 | 1.3861 | 1.4038 | 1.4114 | 1.4095 |
|  | overall | 0.1014 | 0.1336 | 0.2548 | 0.5041 | 0.7581 | 1.2433 | 1.2586 | 1.2697 | 1.2705 | 1.2744 |

In Table 2, we furthermore included the progression of the training loss, since it is an early indicator for model performance on our validation set. Please note that the loss for implicit and explicit learning are based on two different loss functions, refer to Section 2, which explains the difference in magnitude. Moreover, it is also worthwhile looking at the temporal evolution of the same-class and different-class distances, since the structure of the embedded space ultimately influences clustering performance.

Figure 6 visualizes model performance over the time span of 5 epochs for a fixed number of 10^4^ ‘pseudo’-classes, each containing 6 samples only. The plot shows the learning improvement as an increasing amount of images are processed by our proposed deep neural network, introduced in Section 2. We compare the discussed implicit and explicit learning approach by juxtaposing their average overall accuracy as well as average overall distance. In general, we expect both measures to increase over time as more images were seen by the model and utilized to fine-tune parameters.

**Figure 6:**
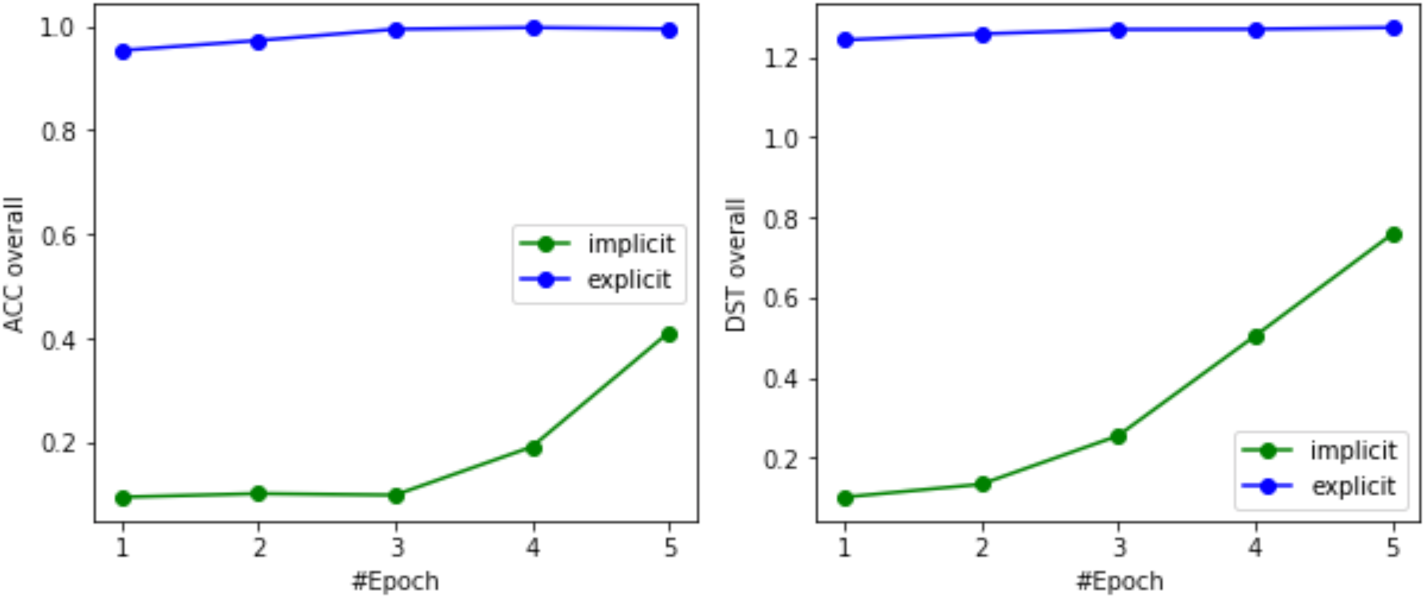
Visual comparison of implicit (green) and explicit (blue) learning approach, showing overall accuracy (left) and overall distance (right) on MNIST benchmark for varying number of epochs/batches, with fixed number of 10^4^ ‘pseudo’-classes, each containing 6 samples only, see Table 2.

#### 4.1.2 Real-World Validation

In case of our real-world biomedical dataset, as described in Section 3.2, the metadata provides information about 57 compound-concentrations or rather ‘pseudo’-classes, which we are utilizing to recover groups of different phenotypes. Since the number of ‘pseudo’-classes is fixed, we just evaluate model performance for a varying number of training epochs/batches. In our real-world experiments, we set the batch size to 11 * 2 = 22 samples, since we assume 11 super-classes and the ‘n-let’ batch construction samples one pair (*2) for each potential super-class, see explanation in Section 2.2.

Our preprocessed BBBC021 training set consists of 2719 × 22 = 59, 818 images of cropped cells, refer to Section 3.2. Our implicit learning model is trained over 5 epochs on 82% of the 59,818 instances, adding up to roughly 5 × 0.82 × 59, 818 ≈ 245, 300 images that were utilized to tune the parameters of our deep neural network. Given 57 ‘pseudo’-classes and assuming about 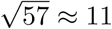 super-classes (or potential phenotypes), we can make a fair performance comparison with our explicit learning model by constructing 245, 300*/*(11 * 2) = 11, 150 batches with 11 image pairs (*2).

It is worth mentioning that we train on cell-level, but validate on image-level. More precisely, for each image we calculate the median of all cell-level embeddings and, subsequently, take the aggregated image-level embeddings to calculate distances, which are in turn used to decide weather two images belong to the same (super-)class or not.

In Table 3 we compare the performance difference between implicit and explicit representation learning for a varying number of training epochs. The following results show that our real-world biomedical dataset is more challenging than the benchmark dataset examined above.

**Table 3:**
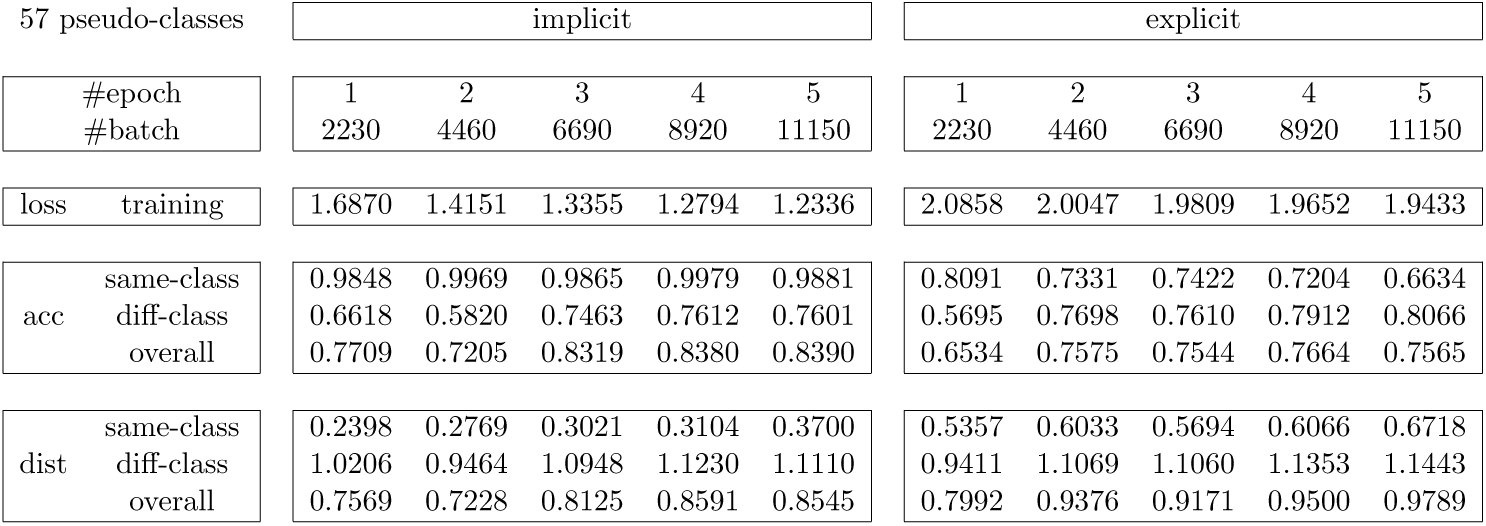
Average accuracy (acc) and average distance (dist) for implicit and explicit learning on prepared real-life BBBC021 dataset (see Section 3.2), for varying number of epochs/batches. The training loss shows how fast our model learns from the processed data and, furthermore, indicates how well the learned embedding is going to group same-class images and separate different-class images for our validation set, as presented in the rows below.

Figure 7 visualizes the temporal evolution of average same-class and different-class distance over the time span of 5 epochs. In general, we want the same-class distance to shrink and different-class distance to grow over time. Furthermore, we want the gap between same-class and different-class distance to be as large as possible. Moreover, the different-class distances should be bigger than one, since this is the default cutoff for deciding if two images belong to same or different classes. These constraints are either implicitly or explicitly formulated in the cost-function used for optimization, refer to Section 2.1 – 2.2.

**Figure 7:**
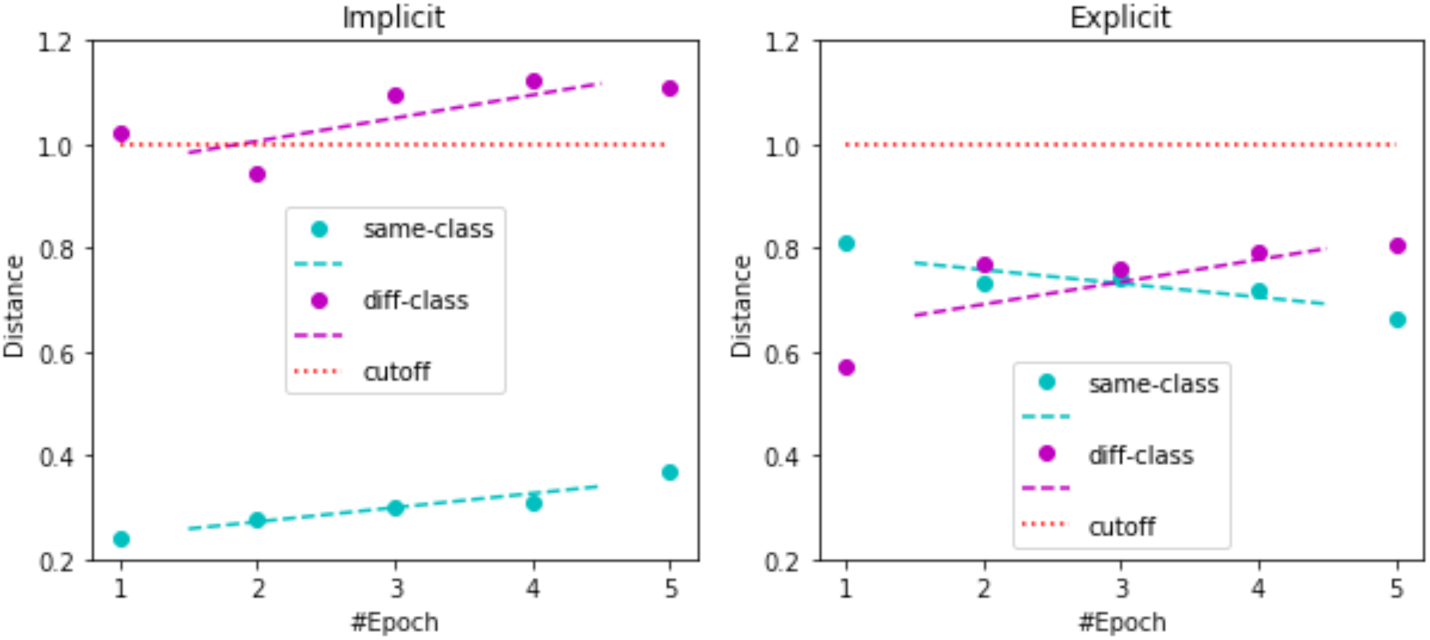
Visual comparison of average same-class (cyan) and different-class (magenta) distances for implicit (left) and explicit (right) learning over time span of 5 epochs, refer to Table 3. We plot linear trend in dashed lines and exact data points as dots. Moreover, we plot the cutoff as a red dotted line.

### 4.2 Cluster Inspection

In this section we take a closer look at the actual cluster formation, resulting from our learned visual representation. Therefore, we fix all parameters of our previously trained model, propagate all validation images through the convolutional neural network, retrieve their lower-dimensional feature representation from the final normalized embedding layer (see Figure 2) and project the feature vectors into a 2D space for visualization, using t-SNE [15] for dimensionality reduction.

#### 4.2.1 Benchmark Inspection

In case of our discussed MNIST benchmark, we decided to plot the cluster formation that results from <u>explicit</u> learning on 10,000 ‘pseudo’-classes after training 1 epoch only, see Table 2. Although this is an extremely challenging task, our explicit learning approach provides a meaningful grouping of the image data, as shown in Figure 8.

**Figure 8:**
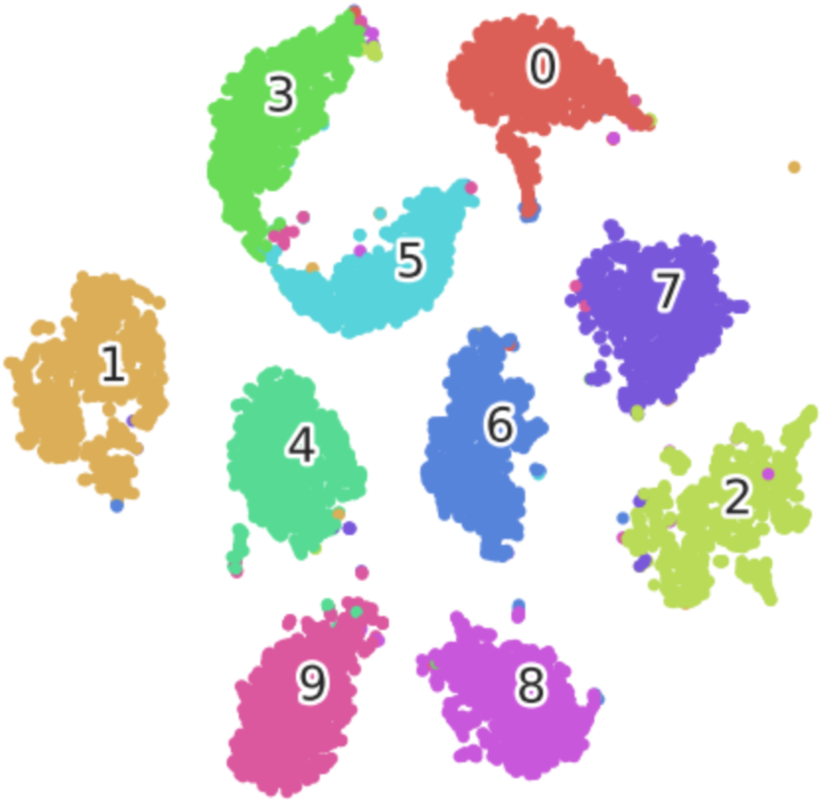
MNIST digits that have been recovered from 10,000 ‘pseudo’-classes by means of explicit representation learning after training for just 1 epoch. Small intra-class distance (0.1355) and high inter-class distance (1.3762) result in an extremely good overall accuracy (0.9569) for grouping same-class and separating different-class images, please refer to Table 2.

Our benchmark cluster inspection (in Figure 8) furthermore illustrates that some digits are easily confused, which is well-known for this dataset. For example, cluster 3 and 5 touch each other, meaning that there are digits that are indistinguishable to our model. However, overall the individual clusters are compact and well separated.

#### 4.2.2 Real-World Inspection

Figure 9 visualizes the cluster formation that was established for the real-world biomedical BBBC021 dataset by means of <u>implicit</u> learning after 5 epochs of training, see accuracy results in Table 3. The plot only shows the embedded images for phenotype ‘DMSO’ and ‘Microtubule Stabalizer’, since these are the two predominating classes with the highest number of tiles, with precise statistics presented at the bottom of the figure.

**Figure 9:**
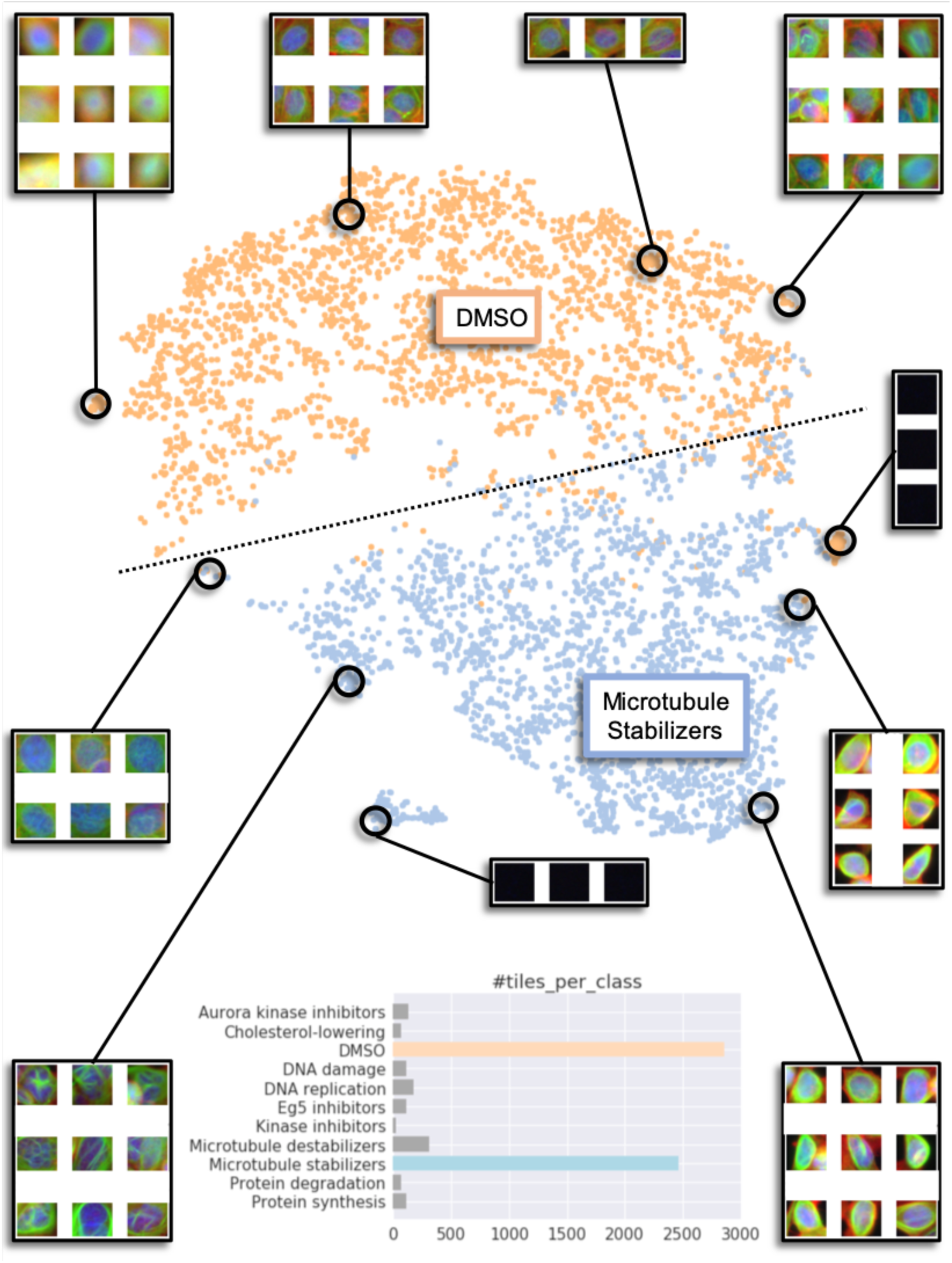
Clustering of the two predominating BBBC021 cell phenotypes that have been recovered from 57 ‘pseudo’-classes/compound-concentrations by means of our implicit representation learning after training 5 epochs. The learned embedding separates the classes well (see detailed accuracy results in Table 3), which is indicated by the horizontal dashed line. Zooming into various cluster regions shows that neighboring cell images exhibit similar visual features and distant cell images display rather different morphological characteristics. Our cluster inspection also uncovered outliers or all black images, which are due to imprecise coordinates used for extracting cells/tiles.

In addition, we zoom into various cluster regions to closely inspect the cell images that were grouped together. The enlarged cluster regions show that neighbouring cell images are not only similar in color and saturation, but also share common morphological characteristics. Furthermore, we can observe that both classes contain visually distinct images, although the intra-class variance is quite high.

Moreover, when looking at Figure 9, we can discern outliers, which are represented as smaller islands of data points at the cluster periphery. In our visual inspection, we can spot images that are all black, since the identified coordinates for cutting individual cells out of the whole microscopy image might have been imprecise. This kind of explorative analysis is extremely valuable, because it can help to uncover bias in measurement equipment, preprocessing steps and modeling techniques.

Overall, our real-world cluster inspection demonstrates that the proposed metadata-guided visual representation learning approach is able to recover and separate individual phenotypes by merely employing information about the applied compound, concentration and internal image structure. The learned embedding can be employed for various down-stream applications, including outlier/novelty detection as well as classification. For example, it is easy to imagine how the two investigated classes could be separated by a simple linear model, as we have indicated by a horizontal dashed line.

## 5 Discussion

Having presented our empirical results for both implicit and explicit visual representation learning in the previous Section 4, we are now in the position to discuss and interpret all tables and graphs.

### 5.1 Benchmark Discussion

As presented in Table 1 and visualized in Figure 5, the performance of our metadata-guided visual representation learning clearly depends on the number of ‘pseudo’-classes. Our results on the prepared MNIST benchmark dataset show that the overall average accuracy of implicit learning drops significantly with an increasing number of ‘pseudo’-classes, whereas explicit learning demonstrates an extremely high and steady performance, even for an incredibly high number of 10,000 ‘pseudo’-classes. This is due to the small intra-class and high inter-class distance achieved by explicit learning, see Table 1, meaning that our <u>explicit</u> constraints on the cost-function have enforced an embedded space with the desired structural properties, grouping same-class and separating different-class images.

Further experiments, summarized in Table 2 and illustrated in Figure 6, reveal that the implicit approach does not only perform better for a high number of ‘pseudo’-classes, but also learns much faster than the explicit approach. For instance, in case of 10^4^ ‘pseudo’-classes, learning explicitly for just 1 epoch leads to higher accuracy (0.9944) than learning implicitly for over 5 epochs (0.4132), see Table 2. This is also reflected in the average different-class distances, which should be above our 1.0000 cutoff in order to allow different-class separation.

Moreover, the training loss, presented in Table 2, indicates how well our model parameters fit the training data. In case of *n* super-classes or rather 10 distinct MNIST digits, the desired loss for explicit learning is give by *loss* = *log*(1 + 9 * *exp*(0.0 – 1.0)) ≈ 1.4611, see Section 2.2, where the average same-class distance is 0.0 and the average different-class distance is 1.0 for all ‘*n*-let’ pairs in a batch. As demonstrated in Table 2, the training loss for explicit learning strategy surpasses our expectation, even for the challenging task of recovering structure from 10^4^ ‘pseudo’-classes. An the other hand, implicit learning stagnates at a cross-entropy loss of ≈ 9.2, which is underwhelming.

In addition, Figure 8 shows that the clustering of our explicit learning approach is meaningful. The plot depicts a clear separation of the MNIST digits within the embedded space. This is quite extraordinary, since the structure was recovered from 10^4^ ‘pseudo’-classes, each containing 6 samples only. This indicates that our metadata-guided visual representation learning is able to actually reveal the hidden structure by pulling together visually similar and pushing away potentially different images.

### 5.2 Real-World Discussion

Our experiments on the BBBC021 dataset show that the grouping of cell images is more intricate than clustering MNIST digits. This is mainly due to the fact that the BBBC021 dataset exhibits high intra-class variance and high inter-class similarity, meaning that we observe (i) heterogeneous cell morphologies in one phenotype and (ii) common visual characteristics among different phenotypes, see Figure 4. Furthermore, the data publishers [3, 13] indicate that only some of the phenotypes were confirmed visually and the remaining were defined based on literature. Since some of the phenotypes are indistinguishable by human eye, we would NOT expect any computer vision algorithm to be able to tell these ‘hypothetical’ classes apart.

Table 3 summarizes our empirical results on the real-world biomedical BBBC021 dataset. We have evaluated the performance of both implicit and explicit learning for a varying number of epochs. In contrast to our previous benchmark, implicit learning is performing better on the cell images than explicit learning. This is primarily due to the small number of 57 ‘pseudo’-classes, which were derived from the individual compound-concentrations found in our prepared dataset. We have already observed this behaviour in our benchmark, see Figure 7, where implicit learning yielded slightly better results for settings with very few ‘pseudo’-classes.

Training over 5 epochs, implicit learning is demonstrating a steady and high same-class accuracy (≥0.98) as well as a continuously growing performance for different-class accuracy (up to ≥0.83). An overall accuracy of ≥0.85 after 5 training epoch is an exceptionally good result, given the afore mentioned variance and confusion. Furthermore, we need to keep in mind that the phenotypes are recovered from metadata only.

Figure 9 illustrates the cluster formation that was constructed by our proposed metadata-guided visual representation learning. The illustration confirms that cell images in adjacent regions of our learned feature space share similar visual appearance, whereas distant data points represent cell images with rather different morphological characteristics.

Hence, we consider our proposed metadata-guided visual representation learning as a valid and sound approach that allows us to process and analyse unlabelled biomedical image datasets that have previously been untapped. It can be applied to large image datasets, assuming <u>no</u> annotation other then the information contained in the metadata, e.g. compound-concentration or other properties like row/column/plate identifier known from an assay.

### 5.3 General Discussion

In addition to discussing results on our benchmark and real-world biomedical dataset, we want to share some general findings and observations that we have made in our empirical study.

First of all, it is important to mention the the implementation of <u>explicit</u> learning is <u>not deterministic</u>, because ‘n-let’ batch construction is stochatic in nature [22]. This leads to the effect that our deep neural network sees images in random order, which might lead to slightly different performance results when training two identical model architectures, since the randomly sampled batches may pose hard constrains or trivial cases, which in turn influences the learning rate.

Furthermore, for <u>implicit</u> metadata-guided visual representation learning we have observed that the training accuracy, which was not reported above, converges toward the following quotient: *#super-classes/#pseudo-classes*. For instance, in case of our MNIST benchmark with 10 ‘super’-classes or digits and 100 ‘pseudo’-classes the training accuracy is close to 10*/*100 = 0.1 after 5 epochs. In other words, for each super-class the corresponding 10 ‘pseudo’-classes are confused 9 out of 10 times, which means that they are falsely assigned to one of the other 9 ‘pseudo’-classes that actually belong to the same ‘super’-class. In that sense, the training loss can give us a first hint on how many ‘super’-classes to expect.

Moreover, we want to emphasize that in our empirical study we have used a cutoff of 1.0 for deciding whether an image pair belongs to same or different class. Another threshold, e.g. around 0.6 to 0.8, might lead to much better accuracy results than those presented above. This becomes more obvious when we take a closer look at the same-class and different-class distances reported in Table 1-3. In Figure 5-7 we have plotted the cutoff, making the potential effect of another threshold visible.

## 6 Conclusion and Future Work

In this work we have introduced a metadata-guided visual learning approach, which employs only context information to cluster images in an unsupervised fashion. More precisely, we have juxtaposed implicit and explicit learning techniques to construct an embedded feature space that arranges images according their visual appearance, without supervision and regardless of missing annotation. We have presented empirical results on benchmark and real-world biomedical images, under different conditions and with various settings. Meaningful clustering results and high classification accuracy both demonstrate the capabilities of our proposed machine learning approach. Hence, we believe that metadata-guided learning has great potential for high content screening and other imaging applications.

In future work we plan to apply our proposed metadata-guided visual representation learning to large-scale high-content-screening assays to automatically identify cell phenotypes and corresponding biological mechanisms of action, without relying on manual annotation. Furthermore, we are going to refine the loss function for explicit learning by employing domain-specific learning constraints that ensure more robust embeddings.

## Supporting information

Code for: (i) dataset preparation, (ii) implicit learning and (iii) explicit learning.

