## Supplementary material for "Metadata-Guided Visual Representation Learning for Biomedical Images": Code for: (i) dataset preparation, (ii) implicit learning and (iii) explicit learning.: mnist3d_explicit.ipynb

```
# %% [markdown]
# Self-Supervised Learning using Explicit Embedding

This work is based on the following paper: https://papers.nips.cc/paper/6200-improved-deep-metric-learning-with-multi-class-n-pair-loss-objective

> "Deep metric learning has gained much popularity in recent years, following the success of deep learning. However, existing frameworks of deep metric learning based on contrastive loss and triplet loss often suffer from slow convergence, partially because they employ only one negative example while not interacting with the other negative classes in each update. In this work, we address this problem with a new metric learning objective called multi-class N-pair loss."

# %% [markdown]
<img src="Nlet01.png" width="1000">
<img src="Nlet02.png" width="1000">

# %% [markdown]
## Preliminaries

# %% [code]
import os
import time
import numpy as np
import numpy.random as rng

import tensorflow as tf

import keras
from keras import backend as K
from keras import optimizers
from keras.models import Model
from keras.layers import Input, Dense, GlobalAveragePooling2D, Lambda
from keras.applications.vgg16 import VGG16
from keras.utils.np_utils import to_categorical

from keras.preprocessing.image import ImageDataGenerator

# settings
os.environ['TF_CPP_MIN_LOG_LEVEL'] = '2'

# %% [markdown]
## Preprocessing

# %% [markdown]
### Load data (MNIST that has been re-shaped to fit VGG16 DL architecture)

# %% [code]
x_train = np.load('/home/spiegst1/EMBED/mnist3d_x_train.npy')
y_train = np.load('/home/spiegst1/EMBED/mnist3d_y_train.npy')
x_valid = np.load('/home/spiegst1/EMBED/mnist3d_x_valid.npy')
y_valid = np.load('/home/spiegst1/EMBED/mnist3d_y_valid.npy')

print('Train Data: ' + str(x_train.shape))
print('Train Label: ' + str(y_train.shape))
print('Valid Data: ' + str(x_valid.shape))
print('Valid Label: ' + str(y_valid.shape))

# %% [markdown]
### Re-label training set (in order to simulate self-supervised learning from meta-data)

# %% [code]
# re-label train
sub = 3
cat = np.argmax(y_train, axis=1) 
srt = np.argsort(np.argsort(cat))
sub = np.tile(range(sub), int(len(cat)/sub))
sub = sub[srt]
subcat = [str(a)+str(b) for a,b in zip(cat,sub)]
_,subcat = np.unique(subcat, return_inverse=True)
y_train = to_categorical(subcat)

print('Train Label: ' + str(y_train.shape))
print('Valid Label: ' + str(y_valid.shape))

# %% [markdown]
## Modeling

# %% [markdown]
### define N-let loss function and batch construction

# %% [code]
class Nlet:
    
    def __init__(self, x_train, y_train):
        self.eps = 1e-7
        self.x_train = x_train
        self.y_train = y_train
        self.nbat = 10 # BE CAREFUL!
        self.ncat = y_train.shape[1]
        self.ndim = x_train.shape[2]
        self.nchn = x_train.shape[3]
        
    def batch(self):
        x_batch = np.zeros((self.nbat*2, self.ndim, self.ndim, self.nchn), dtype=np.float32)
        y_batch = np.zeros((self.nbat*2, ), dtype=np.float32)
        context = rng.choice(range(self.ncat), (self.nbat,), replace=False)
        for cat in range(self.nbat):
            fst, snd = rng.choice([int(e) for e in np.where(self.y_train[:, context[cat]]==1)[0]], (2,), replace=False)
            # tuplet
            x_batch[cat*2+0, :, :, :] = self.x_train[fst, :, :, :]
            y_batch[cat*2+0] = context[cat]
            x_batch[cat*2+1, :, :, :] = self.x_train[snd, :, :, :]
            y_batch[cat*2+1] = context[cat]
        return x_batch, y_batch

    def loss(self, true, pred):
        pred = K.clip(pred, self.eps, 1.0-self.eps)
        lssAll = 0
        for i in range(0, self.nbat*2, 2):
            orgEmb = pred[i+0]
            posEmb = pred[i+1]
            dstOrgPos = K.sqrt(K.sum((orgEmb - posEmb)**2))
            lssOne = 1
            for j in range(0, self.nbat*2, 2):
                negEmb = pred[j+1]
                dstOrgNeg = K.sqrt(K.sum((orgEmb - negEmb)**2))
                if i!=j:
                    lssOne += K.exp(dstOrgPos - dstOrgNeg)
            lssAll += K.log(lssOne) 
        return lssAll/self.nbat
    
nlet = Nlet(x_train, y_train)

# %% [markdown]
### load pre-trained VGG16 model, add embedding layer, re-train top layers to fit our problem (aka transfer learning)

# %% [code]
# trans-fer
trans = VGG16(weights='imagenet', include_top=False)

# added
added = trans.output
added = GlobalAveragePooling2D()(added)
added = Dense(4096, activation='relu')(added)
added = Lambda(lambda x: K.l2_normalize(x,axis=1))(added)

# final
model = Model(inputs=trans.input, outputs=added)

# phase 1-3
for layer in model.layers[:15]:
    layer.trainable = False
for layer in model.layers[15:]:
    layer.trainable = True  
model.compile(optimizer=keras.optimizers.SGD(lr=0.005, momentum=0.9), loss=nlet.loss)

# %% [markdown]
### train with N-let loss on multi-class batches

# %% [code]
# train
mlss = 0.0
for i in np.arange(1,13500+1):
    (x, y) = nlet.batch()
    loss = model.train_on_batch(x, y)
    mlss = mlss + loss
    if(i%2700==0):
        print(str(i) + 'th - Training Batch: ' + str(mlss/i))            

# %% [markdown]
### compare to N-let loss for inter-class distance of 1.0

# %% [code]
# dist
np.log(1+9*np.exp(0.0 - 1.0))

# %% [markdown]
### use embedding layer to project input images into lower-dimensional feature space

# %% [code]
n = x_valid.shape[0] 
m = 4096

x_valid_emb = np.empty([n, m], dtype=np.float32)

for i in range(x_valid.shape[0]):
    x_valid_emb[i, :] = model.predict(x_valid[i:i+1, :, :, :])[0]
    if (i%1000==0):
        print(i)   
        
y_valid_emb = y_valid

print(x_valid_emb.shape)
print(y_valid_emb.shape)        

# %% [markdown]
### define Euclidean distance to measure image dissimilarity in embedded feature space

# %% [code]
def euclid(a, b):
    return sum((a - b)**2)**(0.5)

# %% [markdown]
### evaluate classification performance of embedding (that has been learned from meta-data only)

# %% [code]
enum1 = 0
deno1 = 0
dist1 = 0

enum2 = 0
deno2 = 0
dist2 = 0

num = x_valid_emb.shape[0]
for k in range(num):
    fst, snd = rng.choice(range(0,num), (2,), replace=False)
    dstCat = euclid(y_valid_emb[fst, :], y_valid_emb[snd, :])
    dstEmb = euclid(x_valid_emb[fst, :], x_valid_emb[snd, :])
    if dstCat==0.0:
        deno1 += 1
        dist1 += dstEmb
        if dstEmb<1.0:
            enum1 += 1
    if dstCat>0.0:
        deno2 += 1
        dist2 += dstEmb
        if dstEmb>=1.0:
            enum2 += 1
    
acc1 = enum1/deno1
dst1 = dist1/deno1

acc2 = enum2/deno2
dst2 = dist2/deno2

acc3 = (enum1+enum2)/(deno1+deno2)
dst3 = (dist1+dist2)/(deno1+deno2)

print(acc1)
print(dst1)
print( )
print(acc2)
print(dst2)
print( )
print(acc3)
print(dst3)

# %% [markdown]
## Visualization

# %% [markdown]
### define helper function for plotting results

# %% [code]
%matplotlib inline
import matplotlib.pyplot as plt
import matplotlib.patheffects as PathEffects

import seaborn as sns
sns.set_style('darkgrid')
sns.set_palette('muted')
sns.set_context("notebook", font_scale=1.5, rc={"lines.linewidth": 2.5})

def fashion_scatter(x, colors):
    # choose a color palette with seaborn.
    num_classes = len(np.unique(colors))
    palette = np.array(sns.color_palette("hls", num_classes))

    # create a scatter plot.
    f = plt.figure(figsize=(8, 8))
    ax = plt.subplot(aspect='equal')
    sc = ax.scatter(x[:,0], x[:,1], lw=0, s=40, c=palette[colors.astype(np.int)])
    plt.xlim(-25, 25)
    plt.ylim(-25, 25)
    ax.axis('off')
    ax.axis('tight')

    # add the labels for each digit corresponding to the label
    txts = []

    for i in range(num_classes):

        # Position of each label at median of data points.
        xtext, ytext = np.median(x[colors == i, :], axis=0)
        txt = ax.text(xtext, ytext, str(i), fontsize=24)
        txt.set_path_effects([
            PathEffects.Stroke(linewidth=5, foreground="w"),
            PathEffects.Normal()])
        txts.append(txt)

    return f, ax, sc, txts


# %% [markdown]
### use t-SNE for 2D plot of clusters

# %% [code]
from sklearn.manifold import TSNE

x_valid_tsne = TSNE(random_state=123).fit_transform(x_valid_emb)

# %% [markdown]
### plot clusters that have been recovered from meta-data only

# %% [code]
fashion_scatter(x_valid_tsne, np.argmax(y_valid_emb, axis=1))

# %% [code]
```
