## Supplementary material for "Metadata-Guided Visual Representation Learning for Biomedical Images": Code for: (i) dataset preparation, (ii) implicit learning and (iii) explicit learning.: mnist3d_prepare.ipynb

```
# %% [markdown]
### import tools

# %% [code]
import os
import shutil
import numpy as np

import keras
from keras import backend as K
from keras.datasets import mnist
from keras.utils import to_categorical

from PIL import Image

os.environ['TF_CPP_MIN_LOG_LEVEL'] = '2'

# %% [markdown]
### preprocess data

# %% [code]
# data prep
ndim = 224
ncat = 10
nchn = 3

# data load
(x_trn, y_train), (x_val, y_valid) = mnist.load_data()
nobs_train = y_train.shape[0]
nobs_valid = y_valid.shape[0]

# data reshape
x_train = np.zeros((nobs_train, ndim, ndim, nchn), dtype=np.uint8)
x_valid = np.zeros((nobs_valid, ndim, ndim, nchn), dtype=np.uint8)
for i in range(nobs_train):
    img = Image.fromarray(x_trn[i, :, :])
    img = img.resize((224, 224))
    x_train[i, :, :, 0] = img
    x_train[i, :, :, 1] = img
    x_train[i, :, :, 2] = img
for i in range(nobs_valid):
    img = Image.fromarray(x_val[i, :, :])
    img = img.resize((224, 224))
    x_valid[i, :, :, 0] = img
    x_valid[i, :, :, 1] = img
    x_valid[i, :, :, 2] = img 

# data convert
x_train = x_train.astype('float32')
x_valid = x_valid.astype('float32')
x_train /= 255
x_valid /= 255
y_train = keras.utils.to_categorical(y_train, ncat)
y_valid = keras.utils.to_categorical(y_valid, ncat)    
    
# data dimension
print('Train Data: ' + str(x_train.shape))
print('Train Label: ' + str(y_train.shape))
print('Valid Data: ' + str(x_valid.shape))
print('Valid Label: ' + str(y_valid.shape))

# %% [markdown]
### save data

# %% [code]
np.save('/home/spiegst1/DISSIM/x_train.npy', x_train)
np.save('/home/spiegst1/DISSIM/y_train.npy', y_train)
np.save('/home/spiegst1/DISSIM/x_valid.npy', x_valid)
np.save('/home/spiegst1/DISSIM/y_valid.npy', y_valid)

# %% [markdown]
### load data

# %% [code]
x_train = np.load('/home/spiegst1/EMBED/mnist3d_x_train.npy')
y_train = np.load('/home/spiegst1/EMBED/mnist3d_y_train.npy')
x_valid = np.load('/home/spiegst1/EMBED/mnist3d_x_valid.npy')
y_valid = np.load('/home/spiegst1/EMBED/mnist3d_y_valid.npy')

print('Train Data: ' + str(x_train.shape))
print('Train Label: ' + str(y_train.shape))
print('Valid Data: ' + str(x_valid.shape))
print('Valid Label: ' + str(y_valid.shape))

# %% [markdown]
### plot data

# %% [code]
%matplotlib inline
import matplotlib.pyplot as plt
import numpy.random as rng

ncat = 10
ntil = 20

fig,cell = plt.subplots(ncat, ntil, sharex='col', sharey='row', figsize=(ntil,ncat))

for cat in range(ncat):
    idx = rng.choice([int(e) for e in np.where(y_valid[:, cat]==1)[0]], (ntil,), replace=False)
    for til in range(ntil):
        cell[cat,til].imshow(x_valid[idx[til], :, :, :])

# %% [code]
```
